## Supplementary figures and images for "Uptake of exogenous serine is important to maintain sphingolipid homeostasis in *Saccharomyces cerevisiae*"

### S1 Figure

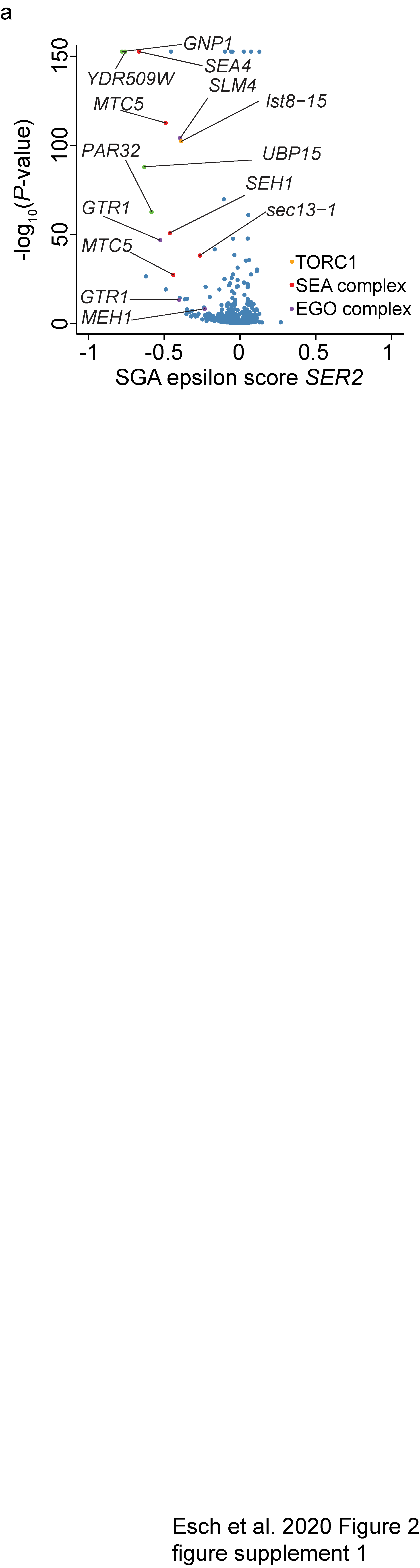

### S5 Figure

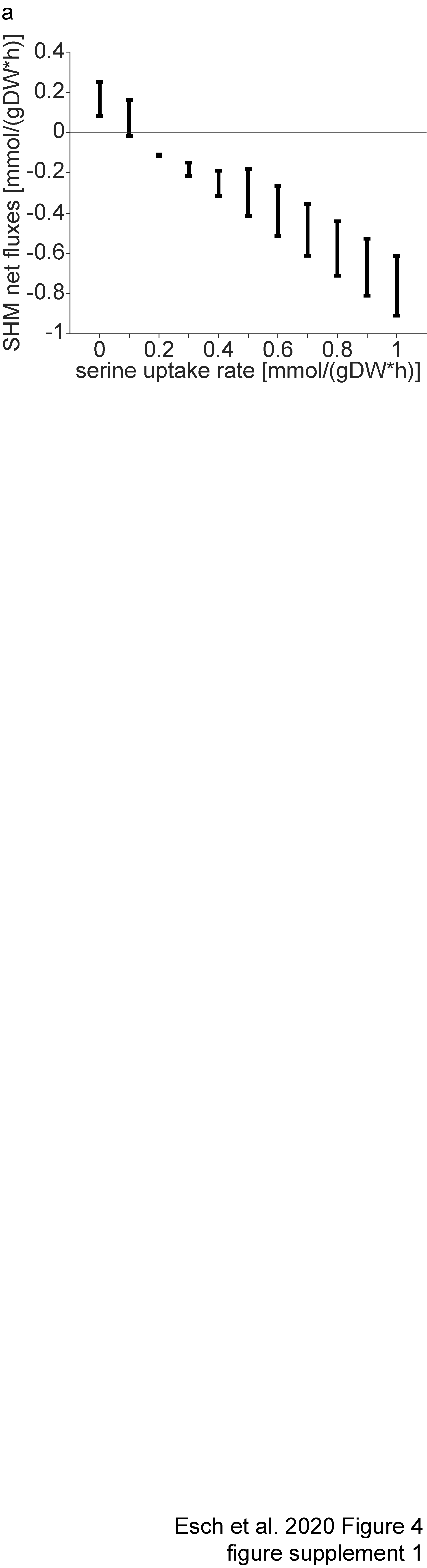

### S7 Figure

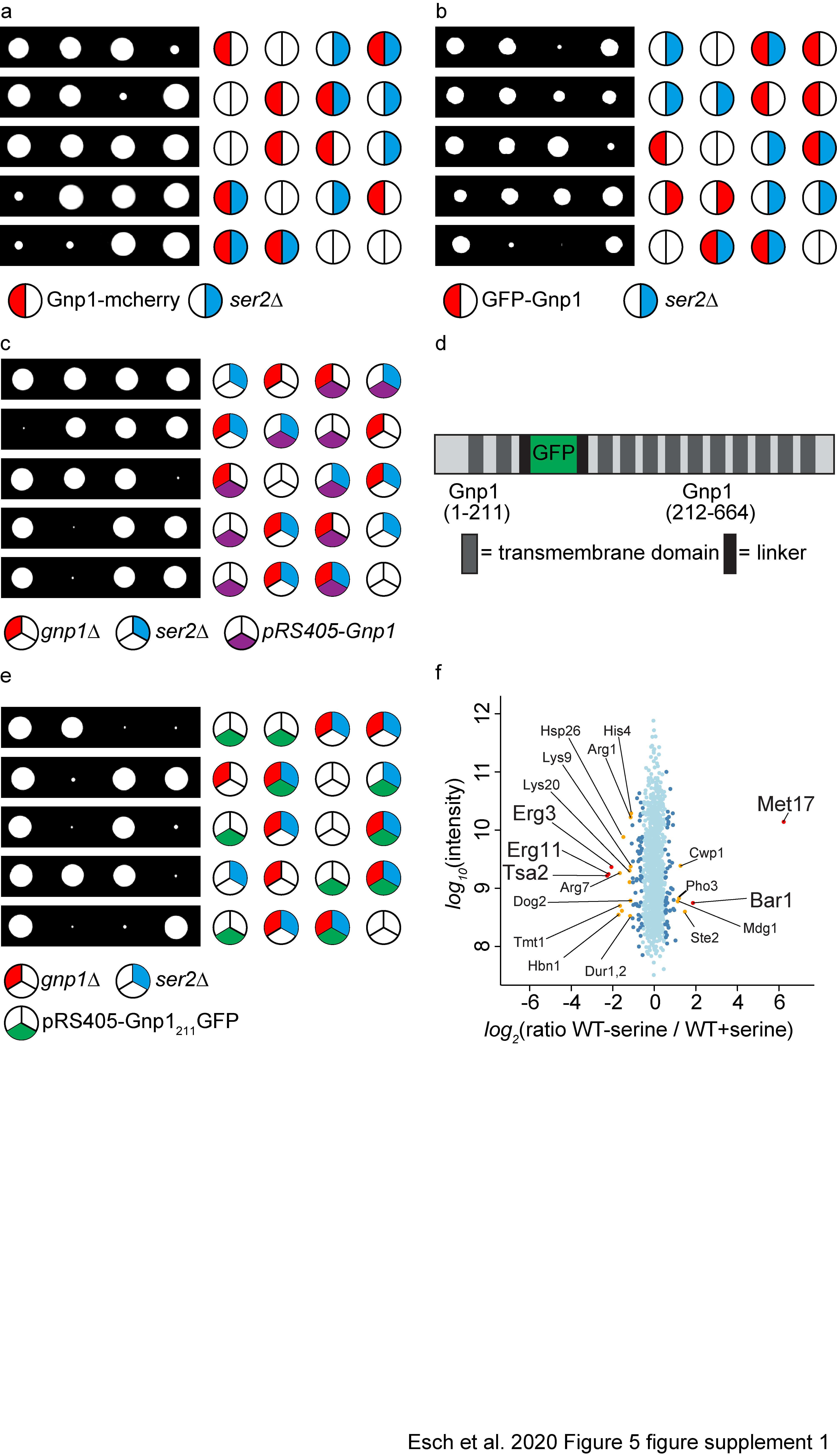

### S8 Figure

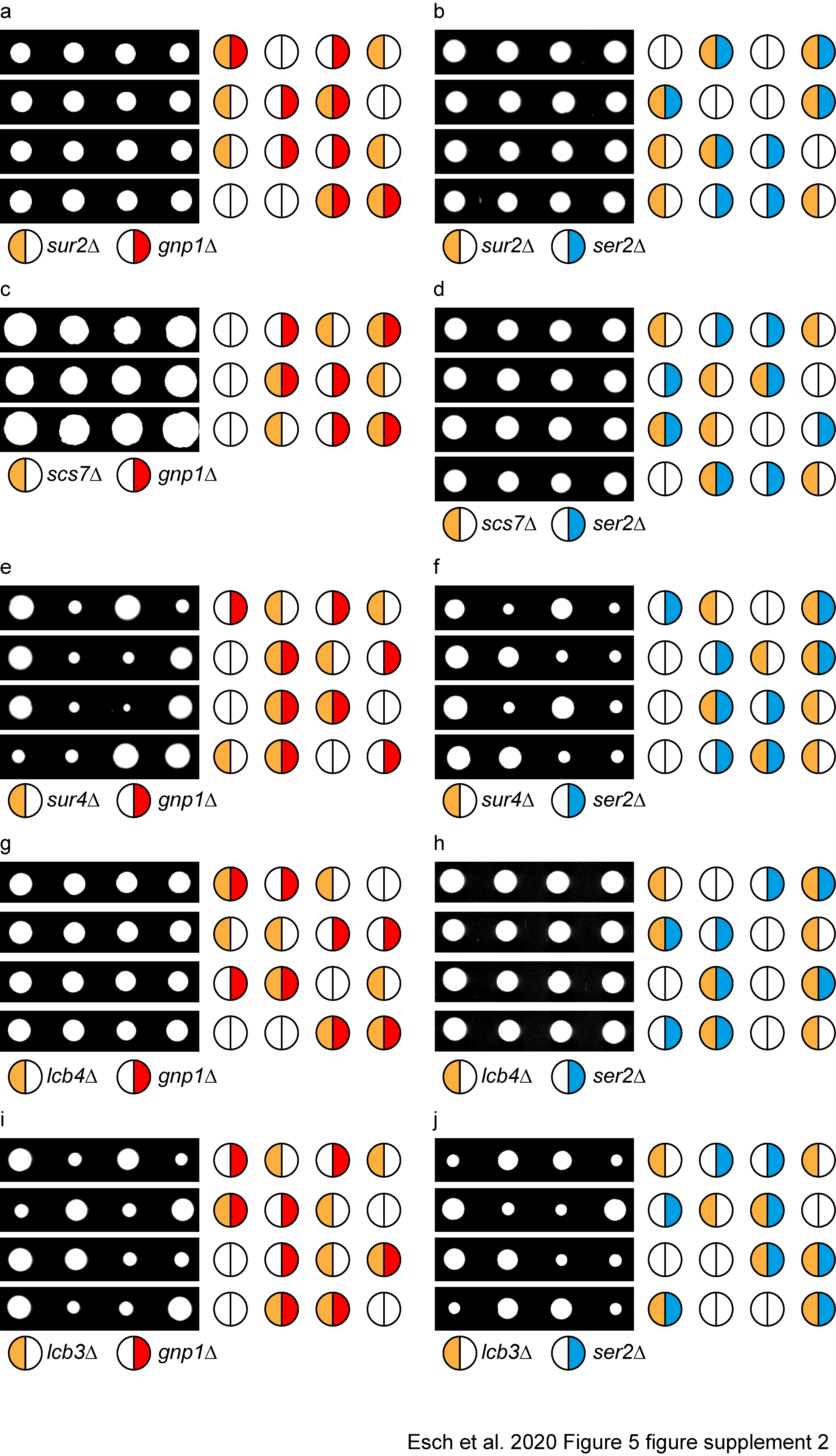

### S9 Figure

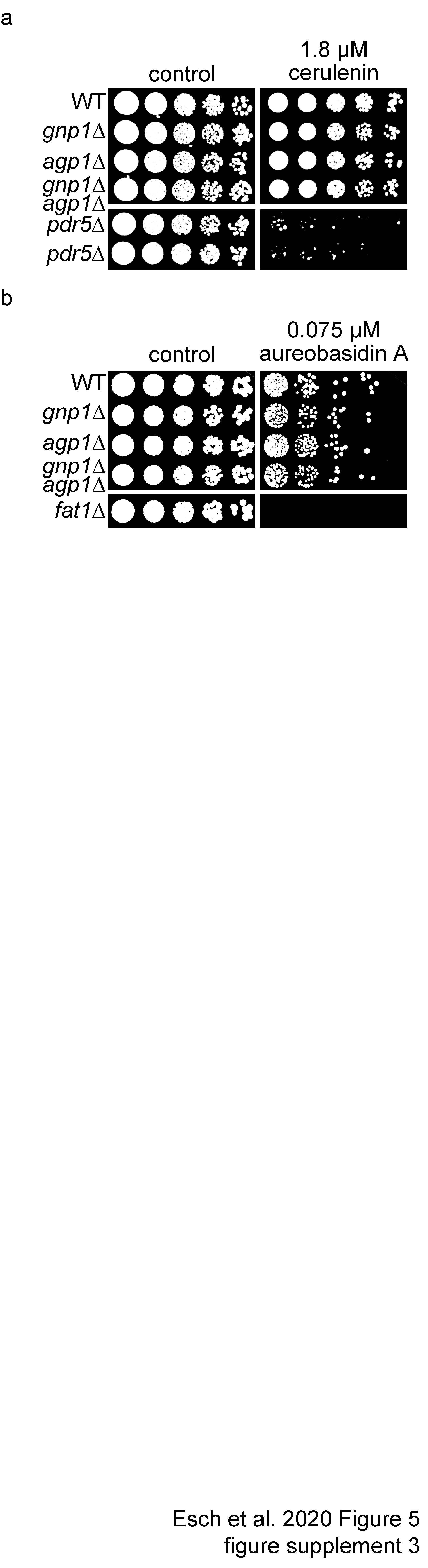

### S13 Figure

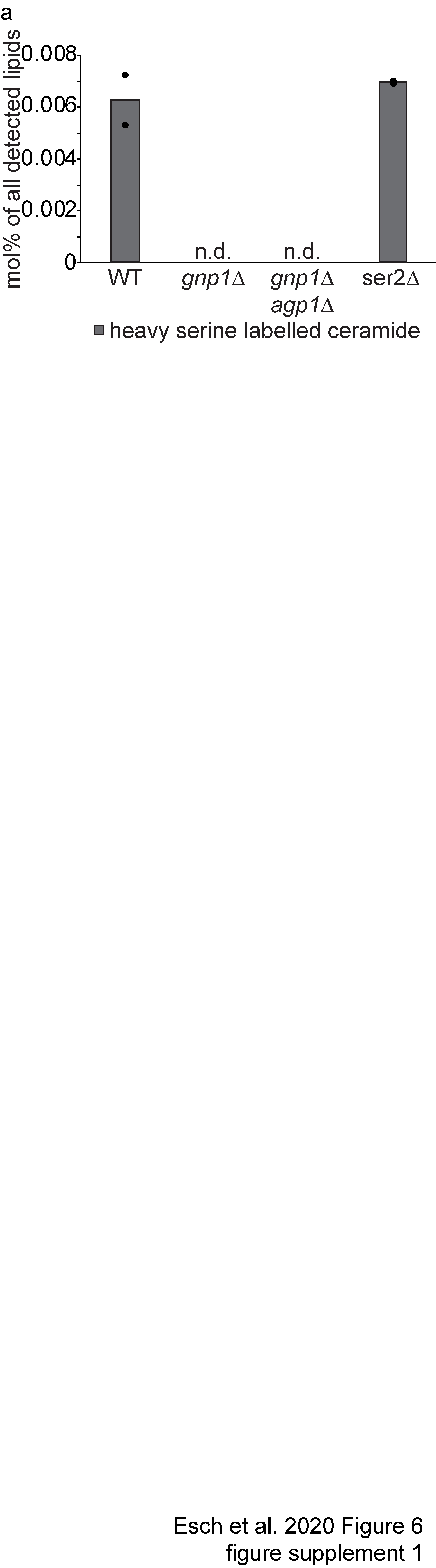
